# Quantitative characterization of the path of glucose diffusion facilitated by human glucose transporter 1

**DOI:** 10.1101/787259

**Authors:** Liao Y. Chen

## Abstract

Glucose transporter GLUT1 is ubiquitously expressed in the human body from the red cells to the blood-brain barrier to the skeletal muscles. It is physiologically relevant to understand how GLUT1 facilitates diffusion of glucose across the cell membrane. It is also pathologically relevant because GLUT1 deficiency causes neurological disorders and anemia and because GLUT1 overexpression fuels the abnormal growth of cancer cells. This article presents a quantitative investigation of GLUT1 based on all-atom molecular-dynamics (MD) simulations of the transporter embedded in lipid bilayers of asymmetric inner-and-outer-leaflet lipid compositions, subject to asymmetric intra-and-extra-cellular environments. This is in contrast with the current literature of MD studies that have not considered both of the aforementioned asymmetries of the cell membrane. The equilibrium (unbiased) dynamics of GLUT1 shows that it can facilitate glucose diffusion across the cell membrane without undergoing large-scale conformational motions. The Gibbs free-energy profile, which is still lacking in the current literature of GLUT1, quantitatively characterizes the diffusion path of glucose from the periplasm, through an extracellular gate of GLUT1, on to the binding site, and off to the cytoplasm. This transport mechanism is validated by the experimental data that GLUT1 has low water-permeability, uptake-efflux symmetry, and 10 kcal/mol Arrhenius activation barrier around 37°C.

**GRAPHICAL ABSTRACT (or TABLE OF CONTENTS ENTRY):** 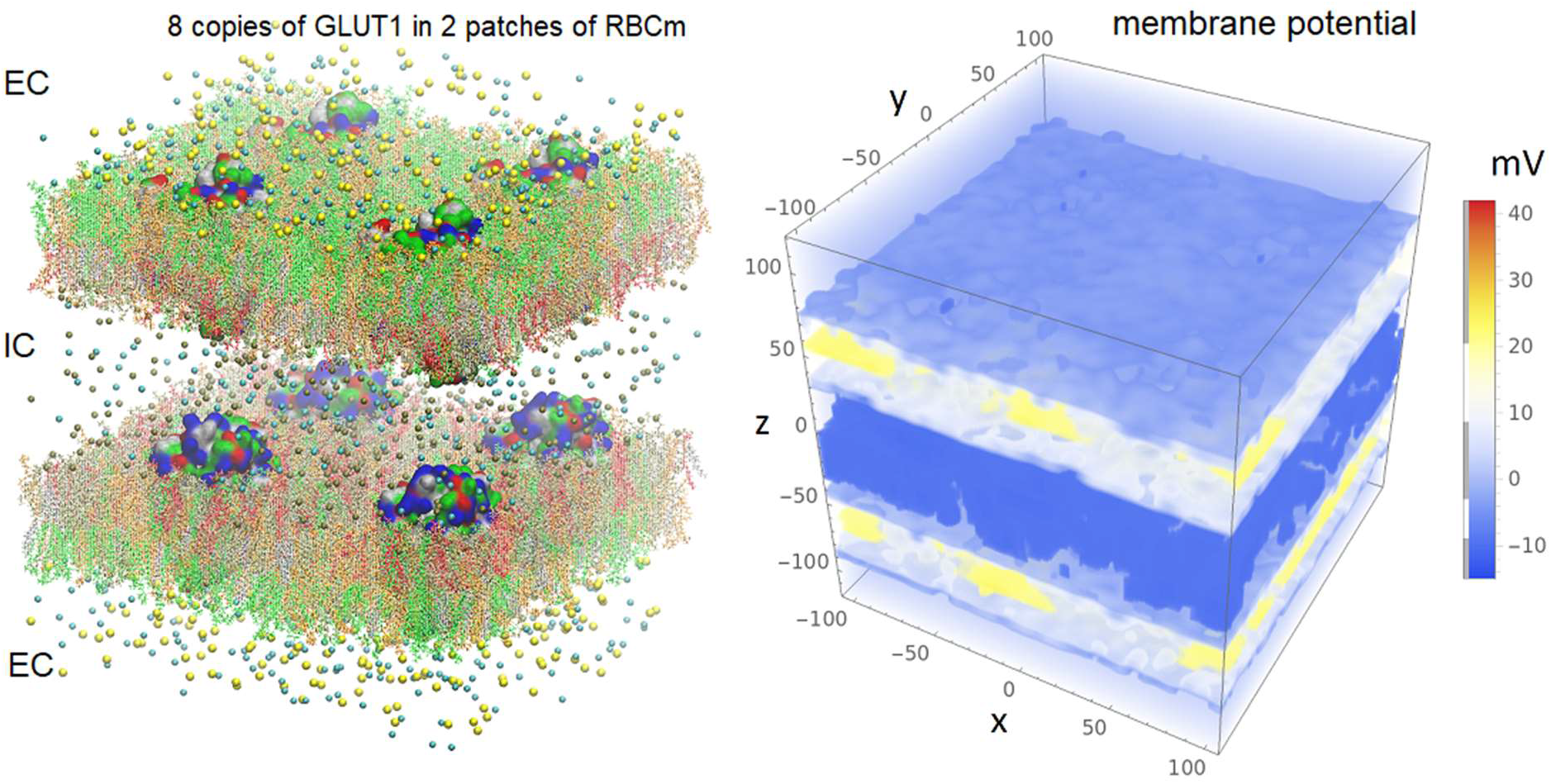

## INTRODUCTION

Thirteen isoforms of glucose transporters (GLUTs)[1-4] are expressed throughout the human body to facilitate a major form of energy metabolism[5-17]. Deficiency in or mutations of GLUTs were found to cause severe health problems[5, 18-25]. In particular, GLUT1[26-29], encoded by *SLC2A1*, is richly expressed in endothelial cells of the blood–brain barrier[7, 17, 30-32]. Deficiency in GLUT1 causes neurological disorders, for example. GLUT1 is replete in erythrocytes[27, 29, 33] and, in fact, it outnumbers water channel protein AQP1[34] that is also very richly expressed in human erythrocytes[33] for hydrohomeostasis. Naturally, GLUT1 has been a subject of many investigations due to its physiological and pathological relevance and due to its abundant availability from erythrocytes.

The structure of GLUT1, like other transporters in the major facilitator superfamily (MFS), has been shown to have a conserved core fold of 12 transmembrane (TM) segments organized into two discrete domains, the amino- and carboxy-terminal (N- and C-) domains[29, 35-40]. And, recently, the crystal structures of GLUT1 have been resolved to atomistic accuracy[39, 40]. In the rich literature of kinetics experiments on GLUT1 (reviewed in, *e*.*g*., [41, 42]), it is unambiguous that glucose transport is symmetric between uptake and efflux both of which are very rapid at near-physiological temperatures. The Arrhenius activation barrier of glucose transport is approximately 10 kcal/mol[42]. It is also unambiguous that GLUT1, even though outnumbering AQP1 (the dedicated water channel) by three folds[33], does not contribute significantly to water transport across the erythrocyte membrane[43].

On the theoretical-computational side, the alternating access theory[1, 44] is generally accepted as applicable for all MFS transporters including GLUT1 beyond the cellular specificities of the membrane lipid-transporter protein interactions. However, it has long been known that the structure and activities of GLUTs are sensitive to the membrane lipid compositions[45-49]. In particular, glucose transport across erythrocyte membranes was found, long ago, to be reduced by 75% by exposure to phospholipase A2 which hydrolyzes fatty acyl groups from the sn-2 position of glycerophospholipids[47, 49]. In general, lipid-protein interactions are significant determinants of the membrane protein functions[50-54]. Therefore, it is natural to ask: Do the specific cellular environments matter at all in how (qualitatively, not just quantitatively) an MFS transporter operates to facilitate moving its substrate in and out of the cell?

In light of all these, surveying the current literature of molecular dynamics (MD) simulations of GLUT1, *e*.*g*., [46, 54-58], and some other GLUTs [55-61], one can see the need to further the current MD studies by modelling a cell membrane with two asymmetries between the intracellular (IC) and the extracellular (EC) sides of the membrane: (1) asymmetric lipid compositions of the inner and the outer leaflets and (2) asymmetric saline compositions of the aqueous compartments on the IC and the EC sides of the membrane. A recent MD study[46] included the asymmetry of the first type mimicking the lipid compositions of human erythrocyte. This study showed that the dynamics of GLUT1 embedded in a patch of lipid bilayer resembling the lipid compositions of human erythrocyte membrane[62, 63] (noted as RBCm) is distinct from GLUT1 embedded in a patch of phosphatidylethanolamine lipid bilayer (noted as POPE). During the unbiased equilibrium MD runs, GLUT1 in POPE patch remained in the endofacial conformation, widely open to the IC side (glucose can easily access the binding site deep inside the protein near its center) and totally occluded to the EC side at both human body temperature (37°C) and subphysiological temperature (5°C). In contrast, at the human body temperature, GLUT1 in RBCm patch presented no significant changes in its endofacial conformation (still widely open to the IC side) but a fluctuating gate on the EC side can open up wide enough for glucose to pass through. This EC gate provides a dynamic passageway for glucose diffusion from the EC side into the binding site near the center of the protein from where glucose can dissociate and move to the IC side, which points to a mechanism of glucose transport through GLUT1 by passive diffusion without the large-scale conformational movements required by the alternating access theory. To valid this EC-gating mechanism, quantitative characterization is needed for glucose diffusion along the passageway connecting the EC saline through the EC gate to the binding site and to the IC saline. Also, water transport through GLUT1 has to be quantified because the passageway for the highly hydrophilic glucose must be favorable for water passage as well. If the EC gate were open most of the time, water transport in human erythrocyte would be mostly by GLUT1 instead of AQP1.

In this article, I present MD simulations of GLUT1 with full consideration of both types of the IC-EC asymmetries (illustrated in Fig. 1 and in the supporting material (SM), Figs. S1-S2), aiming to quantitatively characterize the EC-gating mechanism of GLUT1. In this model system mimicking the human erythrocyte, the asymmetry is considered in lipid compositions of the inner and the outer leaflets of the cell membrane. Also considered is the asymmetry in the IC saline consisting mostly of KCl and the EC saline consisting mostly of NaCl. This latter IC-EC asymmetry is preserved in the MD runs with periodic boundary conditions by employing two membrane patches separating the IC saline compartment from the EC saline compartment. With this, the membrane potential can also be maintained and controlled. In fact, the full electrostatics of the model system gave the membrane potential approximately −40 mV which is reasonably close to the physiological conditions of human erythrocyte. From the unbiased equilibrium MD run of GLUT1 in asymmetric IC-EC environments (and three replication runs), GLUT1 was shown to provide a passageway for glucose diffusion across the cell membrane through a fluctuating EC gate while remaining in the endofacial conformation, in qualitative agreement with Ref. [46]. Computing the probability for the EC gate to be open, the contribution to water transport from 170K GLUT1 copies[33] was estimated to be about 5% of that from the 58K AQP1 copies[33] in an erythrocyte. From the steered MD simulations, the Gibbs free-energy profile of glucose transport was computed in quantitative agreement with the experimental data.

**Fig. 1.**
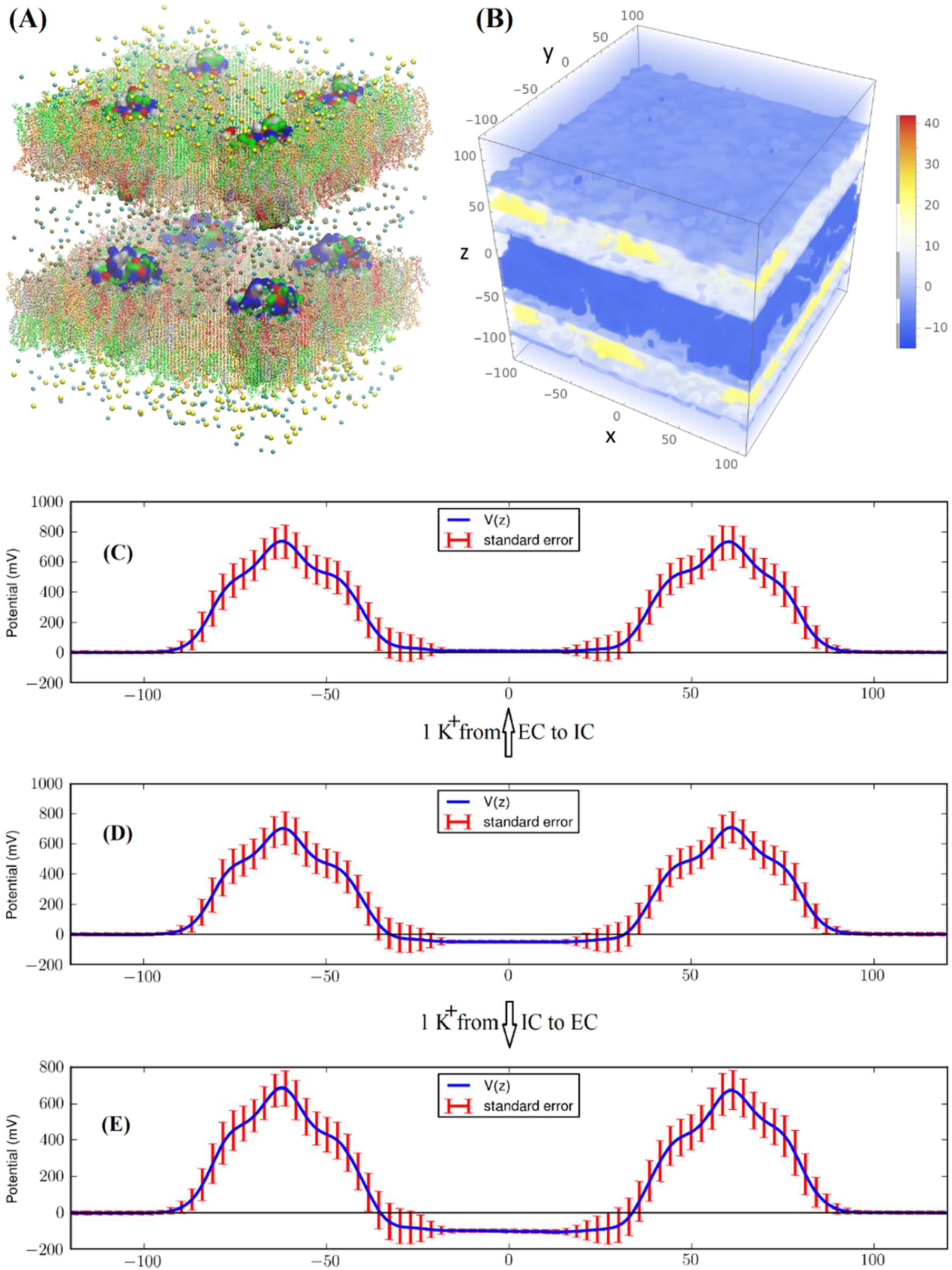
(A) All-atom model system of eight GLUT1 proteins in asymmetric environments mimicking the membrane of a human erythrocyte. The four copies of GLUT1 in the upper patch are numbered as proteins 0, 2, 4, and 6 and the four copies of GLUT1 in the lower patch are numbered as proteins 1, 3, 5, and 7. The system consists of 1,163,104 atoms. The intracellular space is located at −40Å < *z* < 40Å and the extracellular space at *z* < −95Å and *z* > 95Å. The water molecules are not shown for clearer views of all the other constituents of the system. The intracellular salt (KCl) and the extracellular salt (NaCl) are shown as spheres colored by atom names (Cl−, cyan; K^+^, metallic; Na^+^, yellow). The lipids are represented as licorices colored by lipid names (POPC, orange; POPE, tan; POPS, red; SSM, green; CHL, silver). The proteins are presented as surfaces colored by residue types (hydrophilic, green; hydrophobic, white; positively charged, blue; negatively charged, red). (B) The particle mesh Ewald-based electrostatic potential[64, 65]. (C) to (E) The electrostatic potential along the z-axis. The membrane potential of this model system is −40 mV (D), which will decrease to −94 mV (E) (increase to +8 mV (C)) when a single cation is moved across the membrane along the IC-to-EC (EC-to-IC) direction.

## METHODS

The parameters, the coordinates, and the scripts for setting up the model systems, running the simulations, and analyzing the data are available in a dataset at Harvard Dataverse[66].

### Model system setup and simulation parameters

Following the well-tested steps in the literature, CHARMM-GUI[67-69] was employed to build an all-atom model of a single-patch of erythrocyte membrane. CHARMM36 force field parameters[70, 71] were used for the intra- and inter-molecular interactions. NAMD[72] was employed as the MD engine. All simulations were at constant temperature of 37°C and the constant pressure of 1 bar maintained with the Langevin pistons.

I took the high-resolution crystal structure of GLUT1 (PDB code: 4PYP)[39], mutated it back to wild type, removed or replaced β-nonylglucoside with β-D-glucose, translated and rotated the protein complex so that its center is located at the origin of the Cartesian coordinates and its orientation is such that the z-axis points to the IC side, embedded the complex in a patch of lipid bilayer consisting of multiple types of lipids (the IC leaflet consists of 20% cholesterol, 11% POPC, 38% POPE, 22% POPS, and 9% SSM while the EC leaflet has 20% cholesterol, 35% POPC, 10% POPE, and 35% SSM),[62, 63] solvated the sugar-protein-membrane complex with a cubic box of water, and then added sodium/potassium and chloride ions to neutralize the net charges of the system and to salinate the system with 150 mM NaCl. I replicated this single-patch system, inverted the replicate, placed it right next to the original to form a two-patch system, and replaced the IC salt with KCl. Therefore, this all-atom model system of GLUT1 in erythrocyte membrane has an intracellular saline of KCl separated by two membrane patches from the extracellular saline of NaCl. The model membrane has the asymmetry in the lipid compositions of the inner and the outer leaflets mimicking the human erythrocyte membrane. The system so built (consisting of 290,726 atoms) is referred to as SysI. Its dimensions are 108Å × 108Å × 246Å when fully equilibrated. Full details of SysI are shown in SM, Figs. S1 and S2.

### Equilibrium unbiased MD runs of SysI and SysII

After the initial stages of equilibration, I conducted 500 ns production run of unbiased, equilibrium MD on SysI to elucidate the GLUT1 dynamics in the erythrocyte membrane. The particle mesh Ewald (PME) was implemented on a grid level of 128×128×256 for full electrostatic interactions. SysI constituted with two GLUT1 copies in two patches of membrane suffices for proving the concept. Furthermore, taking the coordinates from the last frame of the 500 ns MD run of SysI, I replicated SysI three times. With appropriate translations of the four copies of SysI, I formed SysII illustrated in Fig. 1, a large system of two patches of membrane that consists of 1,163,104 atoms. In each patch of the membrane, there are four GLUT1 proteins. Unbiased MD was run for 500 ns for this large SysII with identical parameters used for SysI except that the PME was implemented on a grid level of 256×256×256. The last 100 ns of the trajectories were used in the computation of the membrane potential (Fig. 1) and the statistics of the EC-gating kinetics.

The large system SysII is advantageous for a very simple reason that the statistics are better with more copies of the transporter protein in a larger system. There are two additional advantages: First, the membrane potential can be fine-tuned in large systems but not in small systems. For example, when a single cation is moved from EC to IC, the membrane potential is altered by a greater amount in a small system (SysI in SM, Fig. S2) than in a large system (SysII in Fig. 1). Second, SysII has a significant enhancement of the signal to noise ratio over SysI because the intrinsic pressure fluctuation of a system is inversely proportional to the volume of the system[73]. These two advantages, while not critical in this study, may be needed in other studies where the biophysical processes are sensitive to the membrane potential or the pressure fluctuations.

### Steered MD for computing the free-energy profile

For the free-energy profile of glucose diffusion through GLUT1, I used SysI with a glucose at the binding site. I followed the multi-sectional approach of [74] to conduct 60 sets of SMD runs to cover the entire diffusion path from the EC to the IC side for a total z-displacement of 60 Å. Over each 1 Å section, the center-of-mass z-coordinate of glucose was pulled/steered at a velocity of 1 Å/ns to sample a forward path and at a velocity of −1 Å/ns to sample a reverse path. At the end of each section, the system was equilibrated for 4 ns while fixing the center-of-mass z-coordinate of glucose. With a total 744 ns SMD simulations, I sampled 4 forward and 4 reverse paths and computed the PMF along the glucose diffusion path, *i*.*e*., the Gibbs free energy of the system when the center-of-mass z-coordinate of glucose is fixed at a given value. The standard errors were over the 4 sets of forward and reverse paths.

## RESULTS AND DISCUSSION

### IC-EC asymmetry and membrane potential

Fig. 1 and SM, Figs. S1-S2 show the model systems for which 500 ns equilibrium (unbiased) MD run was conducted at the physiological temperature 37°C. In order to accurately compute the long-range electrostatic interactions with the PME technique, periodic boundary conditions (PBC) are applied in all three dimensions. The use of PBC leads to artifactitious mixing between the top side with the bottom side of the model system (Fig. 1(A)). In this study, the model system, the unit cell under the PBC, consists of two patches of GLUT1-embedded membranes (Fig. 1) that separate the IC space in between the two patches from the EC space outside the two membranes. The artifactitious mixing is only between the EC sides of the adjacent unit cells. The IC saline and the EC saline are not artifactitiously mixed by the PBC. Any exchange of water or solutes between the IC and the EC can only happen *via* transport across the cell membrane. Consequently, the IC-EC asymmetry is not lost in the PME-based simulations of this work. The IC saline contains 150 mM KCl while the EC saline has 150 mM NaCl. Furthermore, the inner leaflet of the membrane consists of 20% cholesterol, 11% POPC, 38% POPE, 22% POPS, and 9% SSM while the outer leaflet has 20% cholesterol, 35% POPC, 10% POPE, and 35% SSM)[62, 63]. The eight GLUT1 proteins, four on each membrane patch, are oriented in the native orientation as in human erythrocyte.

To illustrate that this model “cell” system is a valid approximation to the human erythrocyte, the charge distribution of the system was computed from the last 100 ns of the 500 ns MD run. The Poisson equation for the electrostatic potential was integrated along the z-axis for the membrane potential shown in Fig. 1. The mean value and the standard error were taken from the statistics over the last 100 ns of the 500 ns MD simulation. The potential difference between the IC side and the EC side of the membrane is approximately −40 mV which is a reasonable approximation for the human erythrocyte. This validates the choice of lipids for the inner and the outer leaflets. Improvement upon this approximation can be furthered with fine tuning the ion compositions of the model system, which is unnecessary in this present study of transport of neutral solutes but will be necessary for any studies of ion transport.

### EC gate fluctuates to open and close without alternating conformational motions

Figs. 2 and 3 representatively show the long-time dynamics of GLUT1 subject to asymmetric IC-EC environments. For both proteins embedded in the two membrane patches of SysI, the dynamics is represented by small fluctuations around the crystal structure. During 500 ns of equilibrium dynamics at 37°C, the protein conformational changes are small and both proteins remain in the endofacial conformation observed in the crystal structure[39, 40] (Fig. 2). The deviations from the crystal structure are mainly from the relatively large sidechain fluctuations and the small shifting of the transmembrane helices. In fact, the deviation as a function of time stabilizes after the initial 100 ns and remains fluctuating under 3 Å (SM, Fig. S3). Interestingly, once the protein dynamics stabilizes at ∼100 ns, a gate on the EC side opens and fluctuates between the open and the closed states. When the EC gate is open, the glucose binding site inside the protein is accessible from the EC side while it is invariably open to the IC. The EC gate, shown in Fig. 3, consists of four groups of residues on four transmembrane helices: (1) Gly31, Val32, Ile33, Asn34, Ala35, and Pro36 (green colored) are located from the EC end of TM1 to the beginning of EC helix 1. (2) Val165, Val166, Gly167, Ile168, Leu169, Ile170, Ala171, Gln172, Val173, and Phe174 (orange colored) on the EC end of TM5. (3) Val290, Phe291, Tyr292, Tyr293, Ser294, Thr295, Ser296, Ile297, Phe298, and Glu299 (violet colored) on the EC end of TM7 and the EC helix connection to TM8. (4) Gln305, Pro306, Val307, Tyr308, Ala309, Thr310, Ile311, Gly312, Ser313, Gly314, Ile315, Val316, and Asn317 (purple colored) on the EC end of TM8. The EC gate, when open, allows glucose to pass through from the EC fluid to the binding site near the center of GLUT1 and *vice versa*. It also allows water permeation through the protein between the EC and the IC sides of the membrane. When closed, the EC gate occludes both glucose and water. Therefore, the probability of the EC gate being open can be estimated from whether or not the EC body of water being connected to the IC body of water as illustrated in Fig. 4.

**Fig. 2.**
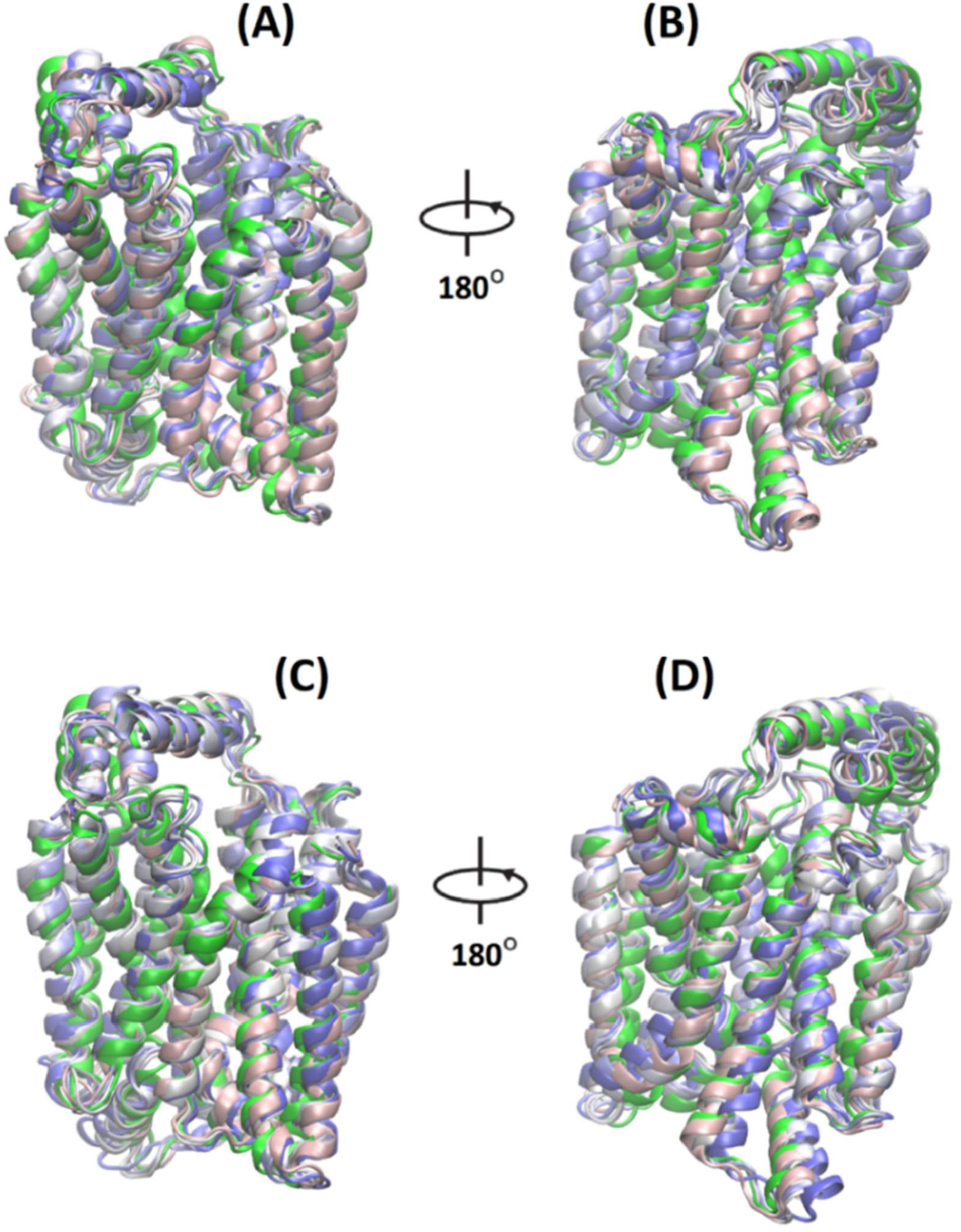
Deviations of GLUT1 from the crystal structure during 500 ns MD run. (A) and (B) show the snapshot structures (grayish blues) of protein 0 at 100, 200, … and 500 ns overlapped on the crystal structure (green). (C) and (D), *i*.*b*.*i*.*d*. for protein 1.

**Fig. 3.**
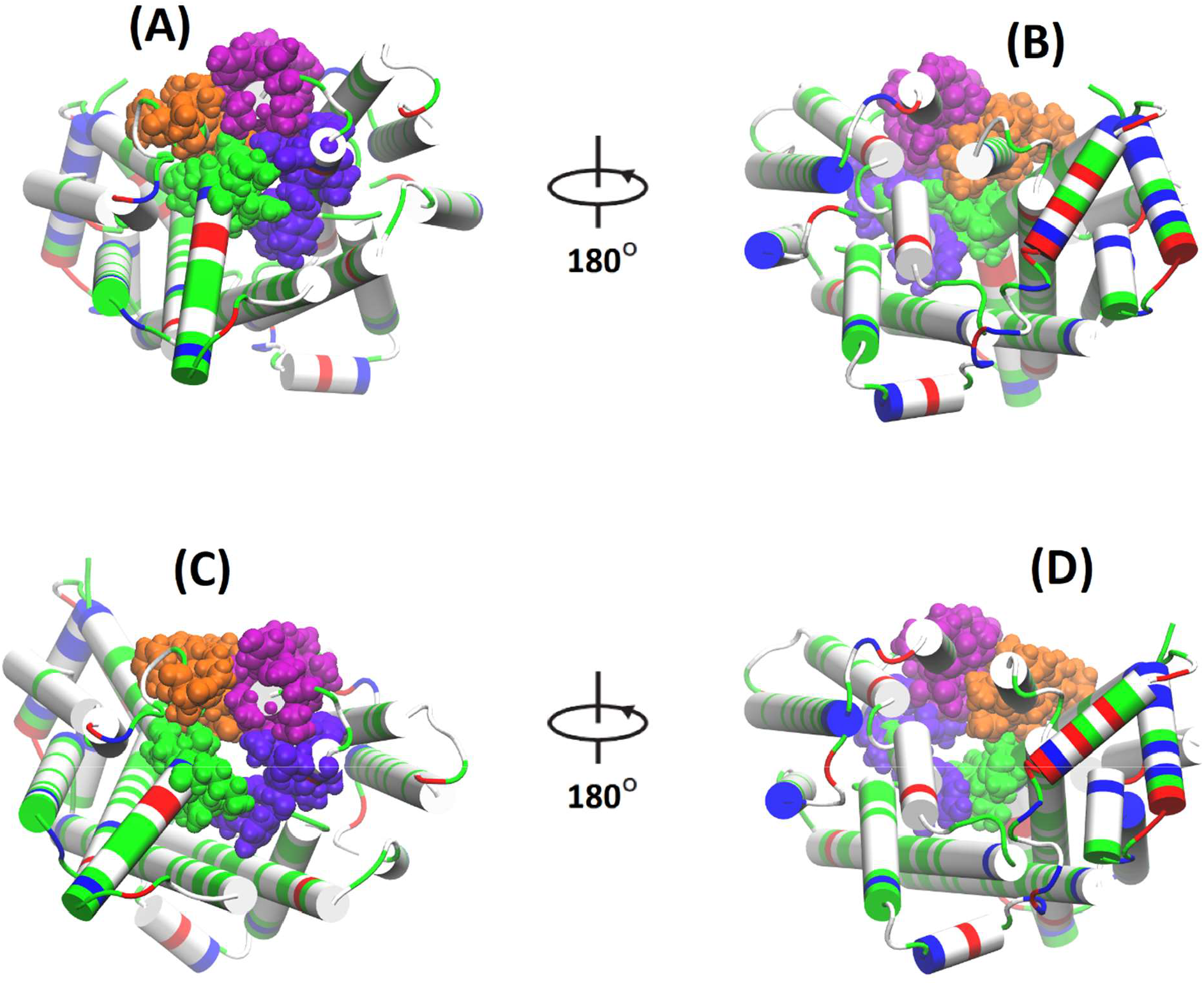
Crystal structure *vs*. dynamic snapshot structure of GLUT1 showing EC-gate closed/open. The EC-gate residues are shown in space-filling spheres colored by the TMs they are on (residues 30 to 38 colored in green, residues 165 to 177 colored in orange, residues 292 to 299 in violet, and residues 305 to 319 in purple) while the entire protein is shown in cartoons colored by residue types (polar, green; hydrophobic, white; acidic, red; basic, blue). (A) and (B) show respectively the EC and the IC views of the protein crystal structure with the EC gate being closed. (C) and (D) show respectively the EC and the IC views of the protein with the EC gate being open (see the space surround by the four groups of the gate residues).

**Fig. 4.**
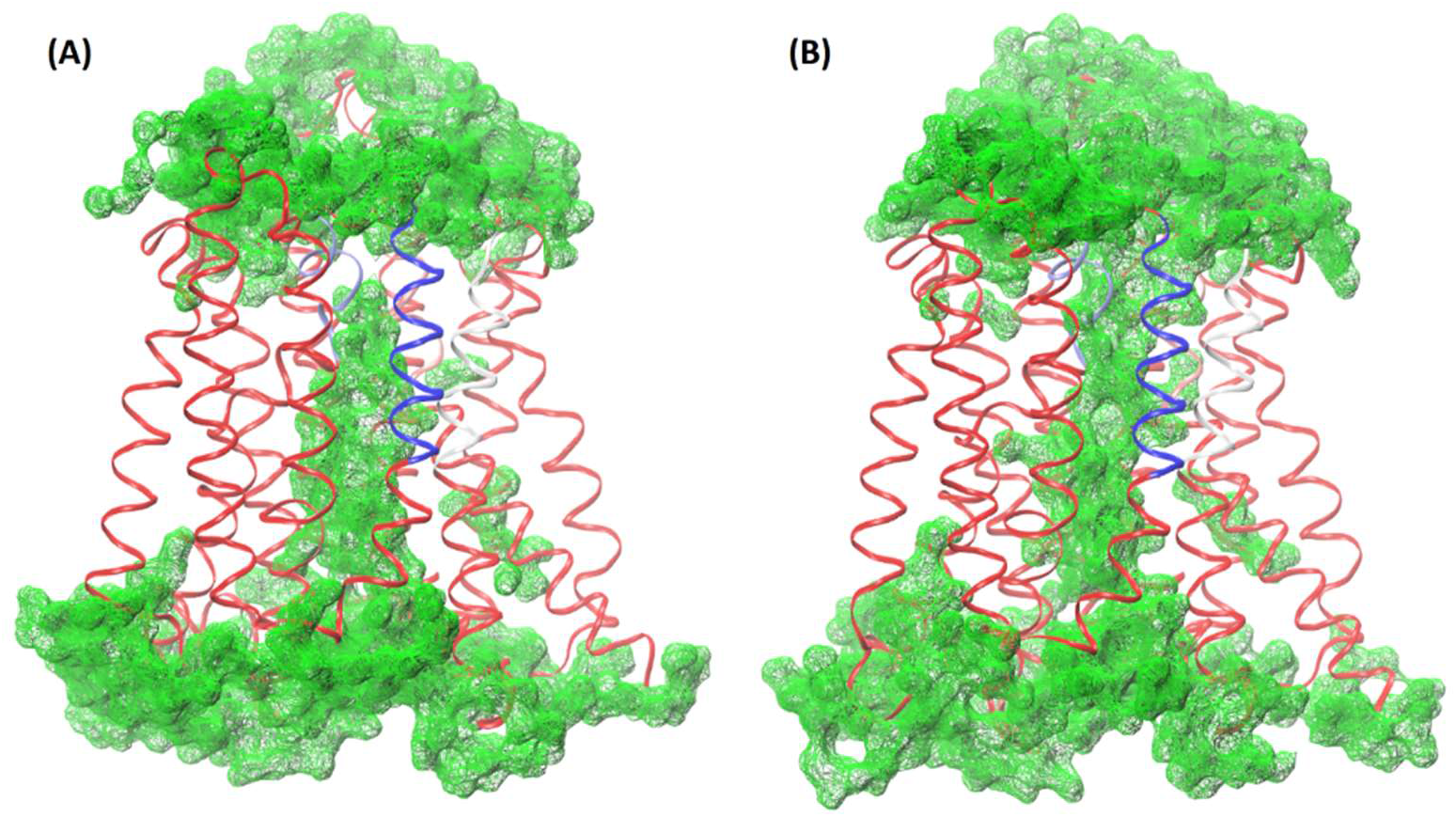
GLUT1 in the EC-gate closed state (A) and in the EC-gate open state (B). The protein is shown in ribbons colored in red except the EC-gate residues. The four groups of the EC-gate residues are colored in blue, white, pink, and cyan respectively. Water molecules within 3.5 Å of GLUT1 are shown in green wireframe surfaces. In the closed state (A), the EC body of water (top) is separated from the IC body of water (bottom). In the open state (B), the two bodies of water are connected (water molecules are within 3.1 Å of each other).

### Statistics of EC gating and water permeability of GLUT1

To achieve reliable statistics of the EC gating mechanism, I conducted 500 ns equilibrium unbiased MD run of SysII that has four copies of GLUT1 (proteins 0, 2, 4, 6) on one patch of the RBCm and four copies of GLUT1 (proteins 1, 3, 5, 7) on the other patch of RBCm as illustrated in Fig. 1. The last 100 ns of the trajectory was used in the statistics of the GLUT1 EC-gating kinetics. The RMSD curves of eight GLUT1 copies are shown in Fig. 5, which shows that none of the eight proteins had large conformational changes during the 500 ns MD run. It should be noted that SysII was formed by replicating SysI in the end of a 500 ns MD run. Additionally, I repeated the simulation of SysII three more times (The RMSD curves are shown in SM, Figs. S4 to S6).

**Fig. 5.**
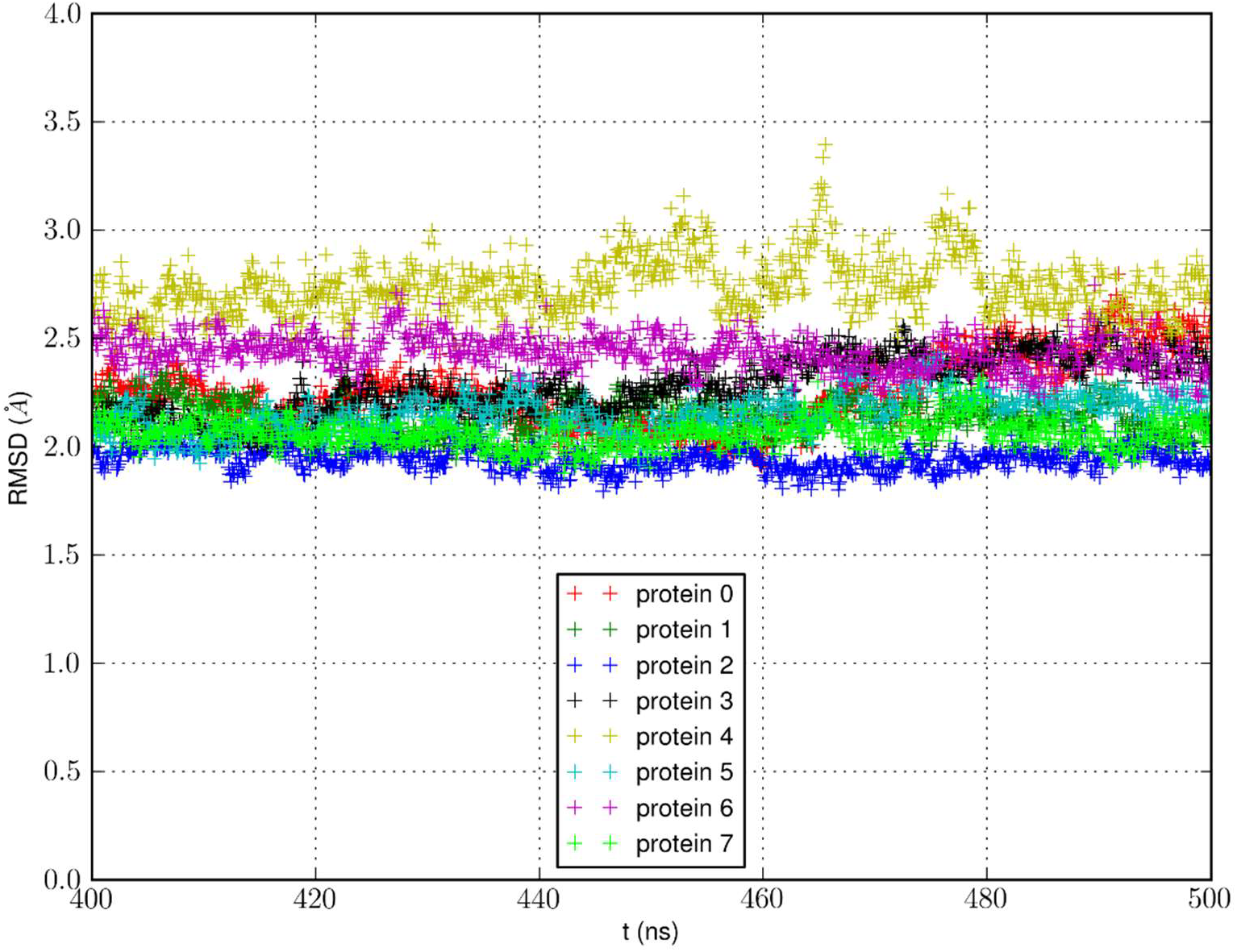
RMSD of eight GLUT1 copies in the erythrocyte membrane from the crystal structure during the last 100 ns of the 500 ns production run of SysII.

Fig. 6 shows the statistical characteristics of the EC gating in each of the eight GLUT1 proteins. The statistical average over the four MD runs gave the probability of the EC gate being open as 0.017 ± 0.005. Assuming that GLUT1 in the EC-gate open state is similar to AQP1 which is always open for water conduction, the 170 K GLUT1 proteins on a human erythrocyte would give a contribution to water transport at ∼5% of the water conduction facilitated by the 58 K AQP1 proteins that are the dedicated water channels of an erythrocyte. Qualitatively, the GLUT1 dynamics simulated in SysI or SysII are similar to the case studied in Ref. [46] where GLUT1 was embedded in a single membrane patch that had similar lipid compositions. Quantitatively, the dynamics of GLUT1 in the fully asymmetric EC-IC environments are substantially different from Ref. [46] where only the asymmetry in lipid compositions was considered. In the larger and more realistic model system of GLUT1 in RBCm studied here, the EC gate is open for ∼1.7% of the time whereas, in the small model system of a single-membrane patch, the EC gate was nearly open all the time at 37°C[46]. If the latter were true, then the 170K GLUT1 copies would dominate the 58K AQP1 copies in water transport across the erythrocyte membrane.

**Fig. 6.**
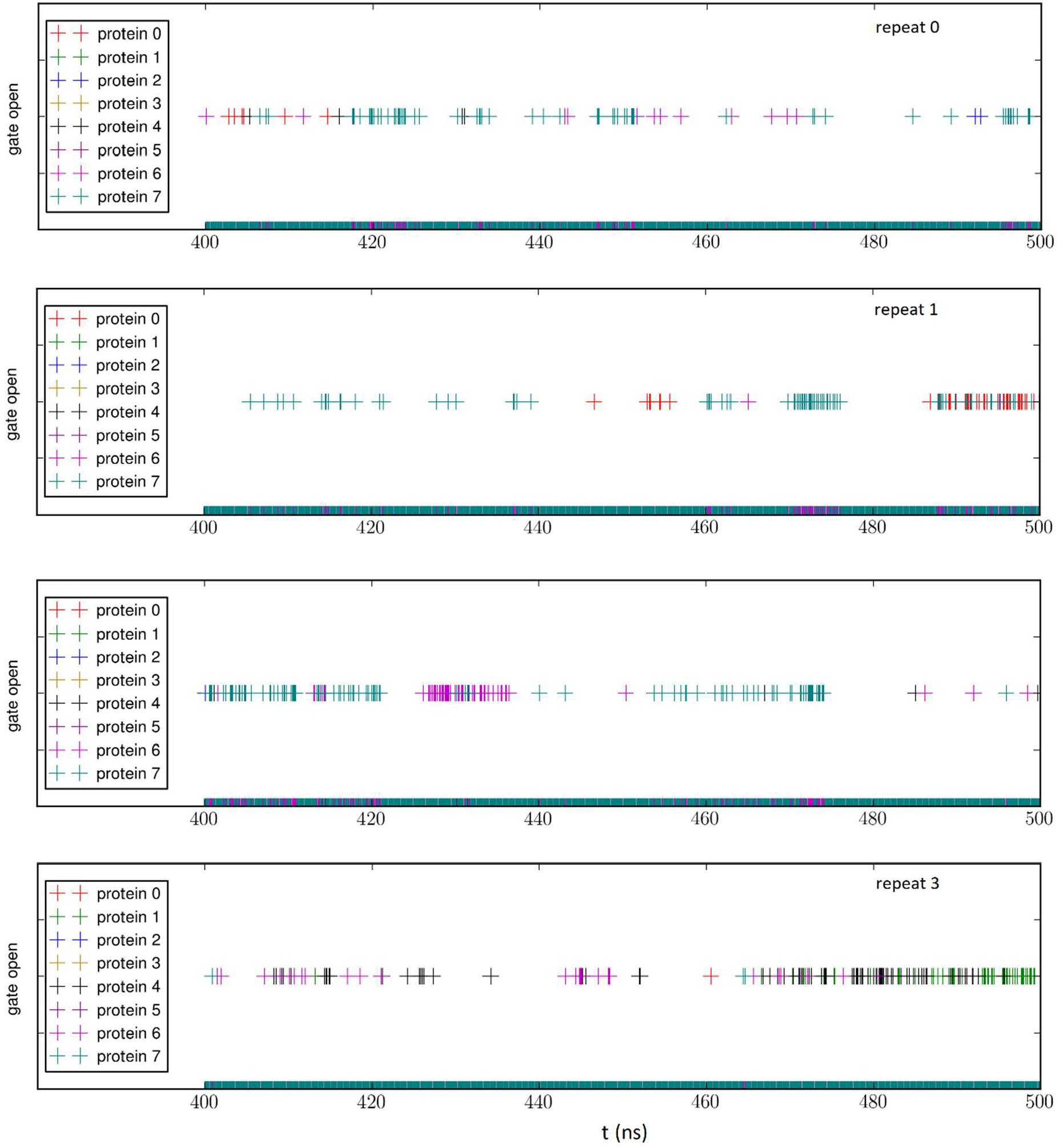
EC-gating kinetics during the last 100 ns of the 500 ns production runs. The criterium for the EC-gate to be open is defined in Fig. 4. The probabilities for the EC gate to be open are, respectively, 0.0113, 0.0147, 0.0205, 0.0207 for the four MD runs of SysII (from top to bottom). Altogether, these four data sets give a mean of 0.0168 with a standard error of 0.0046.

### Quantitative characteristics of glucose transport

Fig. 7 shows the Gibbs free energy along the path of glucose transport through GLUT1. It is known that GLUT1 is a passive facilitator for glucose diffusion down the concentration gradient. The driving force for glucose transport is the Gibbs free-energy gradient along the diffusion path. Experimental data of kinetics showed an Arrhenius activation barrier ∼10 kcal/mol around 37°C.[42] This high barrier renders it infeasible to compute the free-energy profile directly from equilibrium MD simulations. Shown in the top panel of Fig. 7 are the results of 744 ns SMD runs detailed below:

**Fig. 7.**
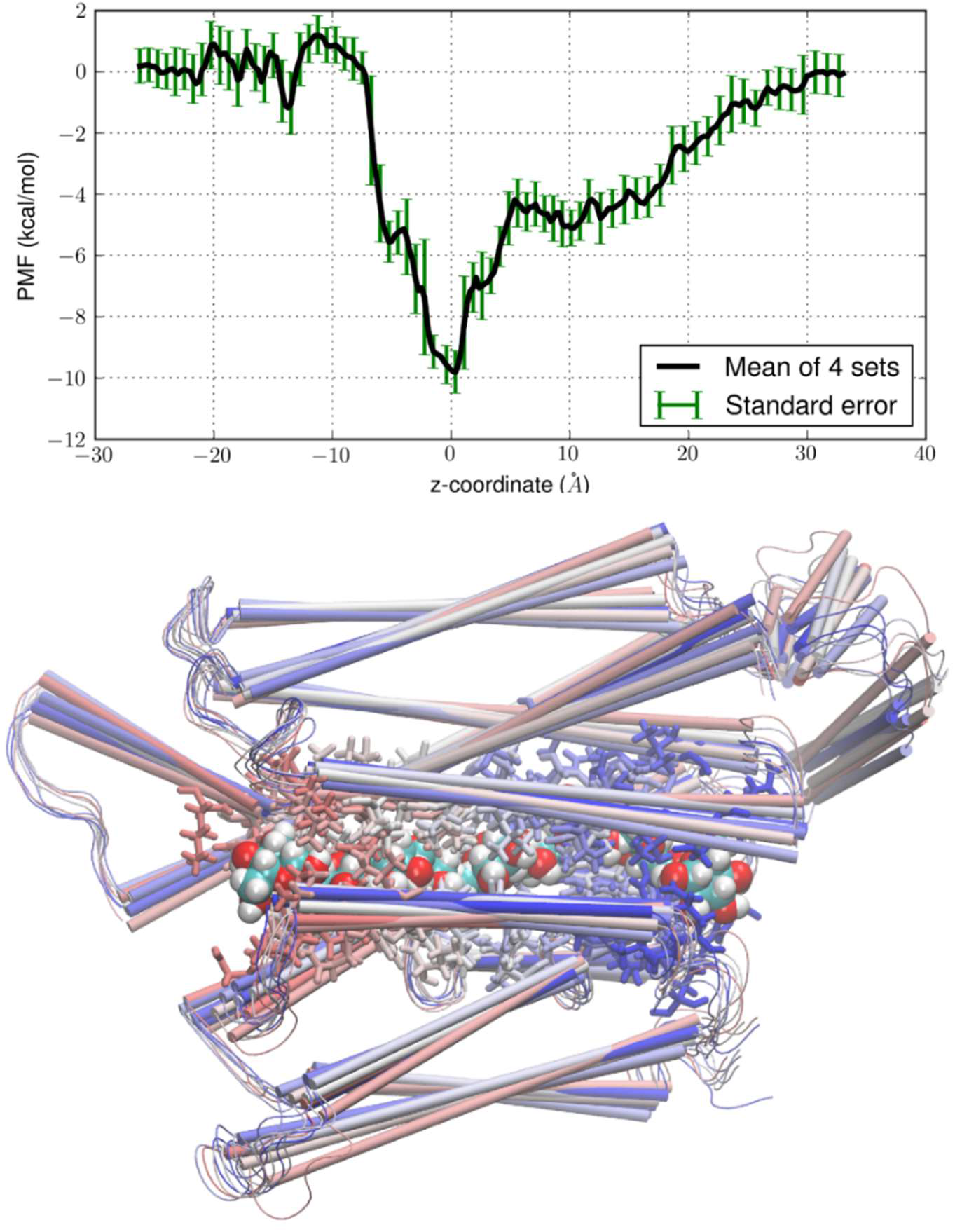
Glucose transport path of diffusion facilitated by GLUT1. Top, PMF of glucose along the diffusion path computed from SMD simulations. The PMF(z) represents the Gibbs free energy of the system when the center-of-mass z-coordinate of glucose is at a given position. Bottom, glucose (space-filling spheres), GLUT1 (thin cartoons), and residues within 5 Å of glucose (licorices) shown in representative frames along the transport path. The sugar is colored in red (O), cyan (C), and white (H) in all frames but the protein is colored by the fame.

Starting from the binding site (*z* = 0 Å), the glucose molecule was pulled/steered forward and backward over sections of 1 Å each: 32 sections to the IC side and 28 sections to the EC side. In each section, the center-of-mass z-coordinate of glucose was steered at a speed of 1 Å/ns along the z-axis in the sampling of a forward path and along the negative z-direction in the sampling of a reverse path. The pulling was repeated 4 times to sample 4 forward and 4 reverse paths in each section. At the end of every section, the system was equilibrated for 4 ns while the z-coordinate of glucose’s center of mass was fixed. The x- and y-coordinates of the glucose center of mass were free to follow the stochastic dynamics of the system.

The work done to the system along the forward paths and that along the reverse paths both contain irreversible dissipative work in addition to the reversible change in the Gibbs free energy. The use of the Brownian dynamics fluctuation-dissipation theorem had those two parts cancelled to yield the reversible potential of mean force (PMF) as a function of z shown in Fig. 7, namely, the Gibbs free energy of the system when the center-of-mass z-coordinate of the glucose is fixed at a given value[74].

Going along the glucose diffusion path from the EC side (*z*∼ − 30 Å), through the EC gate, down to the binding site (*z*∼0 Å) and from there up to the IC side (*z*∼30 Å), the free-energy difference between the IC and the EC is approximately zero. The equal levels on the two sides are required for passive diffusion of neutral solutes down the concentration gradient. Discrepancy from this equality indicates the inaccuracy of a computational work. On the EC side, through the EC gate, the PMF curve exhibits minor dips and bumps, which reflects that the EC gate constantly fluctuates between closed and open states. The EC-gating residues are flexible to allow glucose passage without significant water flux through there. The PMF at the binding site is approximately 10 kcal/mol below the EC or IC level. This agrees with the experimental fact of glucose-GLUT1 affinity. Each glucose transport event, either uptake or efflux, consists of two parts: falling into the binding site with a PMF drop of 10 kcal/mol and climbing out of there with a PMF rise of 10 kcal/mol. This thermal activation nature of glucose transport quantitatively agrees with the experimental data of 10 kcal/mol in Arrhenius activation barrier around 37°C[42]. The lack of significant barriers above the bulk level PMF on either the IC or the EC side corresponds well with the experimentally observed symmetry between the glucose uptake into and efflux out of human erythrocytes.

The residues along the glucose transport path: Phe26, Thr30, Gly31, Asn34, Ala35, Gln37, Lys38, Thr137, Pro141, Arg153, Gly154, Gly157, His160, Gln161, Ile164, Ile168, Gln172, Leu176, Gln282, Gln283, Ile287, Asn288, Phe291, Tyr292, Ser294, Thr295, Val307, Thr310, Glu329, Arg333, Trp388, Phe389, Gly408, Asn411, and Asn415 (illustrated in Fig. 7). It is interesting to note that 30 of these 35 residues are conserved between human GLUT1 and GLUT3. [46] The many aromatic residues along the path provide affinity for glucose and hydrophobicity to disfavor water passage. All these, in agreement with experimental findings, indicate validity of this research.

## CONCLUSIONS

In three aspects, this *in silico* study reached quantitative agreement with the available experimental data on glucose transport across the erythrocyte membrane: (1) GLUT1 does not contribute significantly to water movement across the erythrocyte membrane. Even though GLUT1 outnumbers AQP1 (the water channel) in human erythrocytes by more than two folds, its contribution to water transport is around a few percent of the total water permeability. GLUT1 is only in the EC-gate open state for less 2% of the time. (2) Glucose transport at near physiological temperatures (∼37°C) is approximately symmetric between the uptake and the efflux directions. (3) The Arrhenius activation barrier of glucose transport around 37°C is approximately 10 kcal/mol. All these aspects consistently and quantitatively point to a very simple mechanism of glucose diffusion facilitated by GLUT1: This membrane protein is a channel with a fluctuating gate.

## Supporting material

Three videos and six additional figures that are discussed but not included in the main text.

## Data availability

The Dataset (parameters, coordinates, scripts, *etc*.) to replicate this study is available at Harvard Dataverse[66].

## Author contributions

LYC is responsible for all aspects of this paper.

## Acknowledgements

The author acknowledges the Texas Advanced Computing Center (TACC) at The University of Texas at Austin for providing HPC and visualization resources that have contributed to the research results reported within this paper. URL: http://www.tacc.utexas.edu.

