## Supplemental figures for "Quantitative characterization of the path of glucose diffusion facilitated by human glucose transporter 1"

L. Y. Chen

This supporting material contains three avi videos and five additional figures that are described as follows:

Movies 1 to 3 show the path of glucose uptake viewed from the extracellular, the intracellular, and the membrane sides respectively.

Figures S1 and S2 illustrates the model system details and the membrane potentials of SysI.

Figures S3 illustrates the root mean squared deviations from the crystal of GLUT1 for the two transporter copies in erythrocyte membrane during the 500 ns run of SysI.

Figures S4 to S6 illustrate the root mean squared deviations from the crystal of GLUT1 for the eight transporter copies in erythrocyte membrane during the last 100 ns of the three repeat runs of 500 ns MD simulations of SysII.

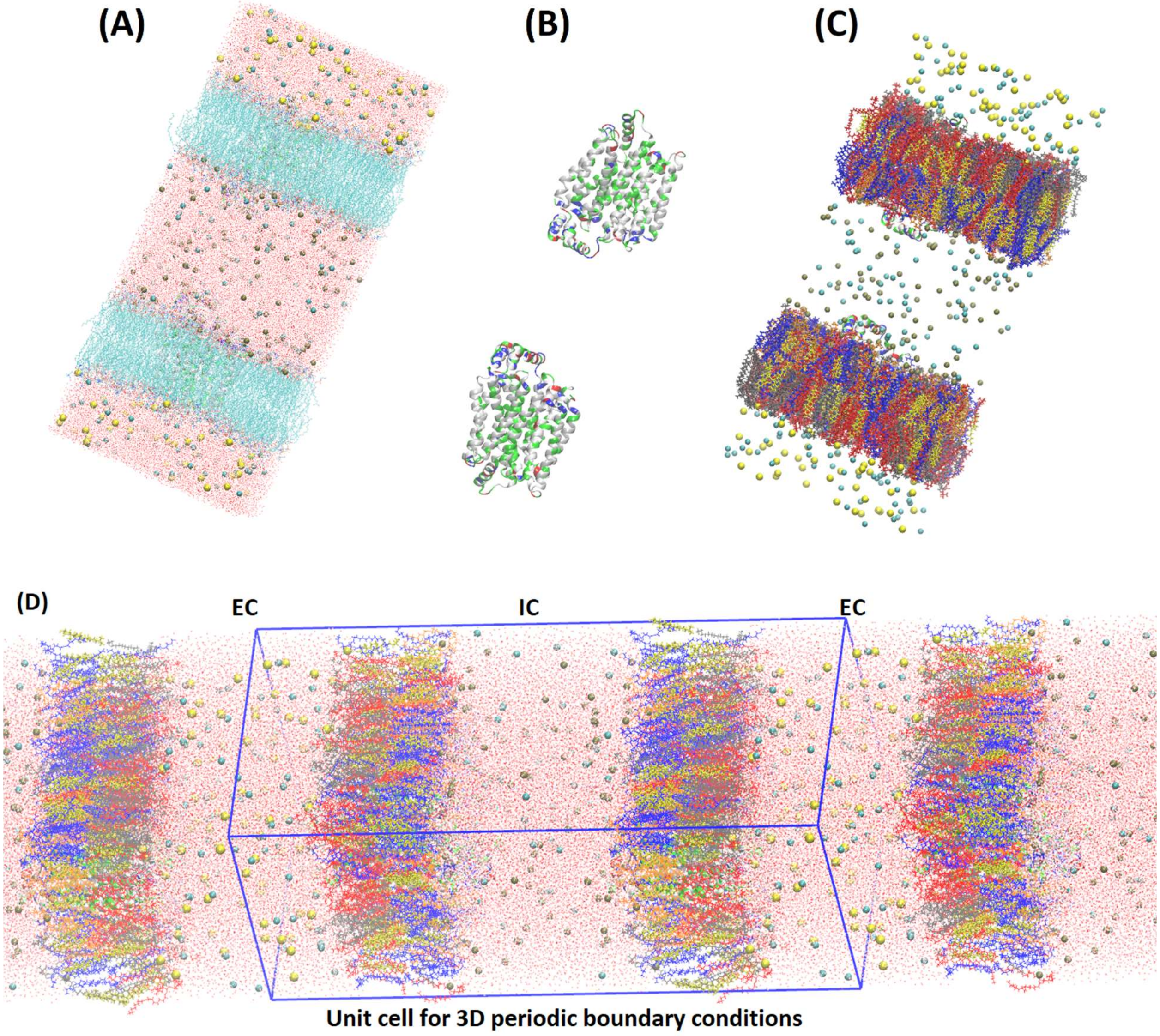


**Fig. S1**. SysI, an all-atom model system of two GLUT1 proteins embedded in two membrane patches separating the intracellular space of KCl saline from the extracellular space of NaCl saline. (A) The entire system consisting of 290,776 atoms. Water and the membranes are shown in lines colored by elements. Ions are shown in large spheres colored by atoms. (Colors by elements/atoms: Na, yellow; K, metallic; Cl, cyan; H, white; C, cyan; N, blue; O, red; P, purple; S, yellow.) (B) Proteins 0 and 1 are shown in ribbons colored by residue types. (Colors by residue types: hydrophilic, green; hydrophobic, white; negatively charged, red; positively charged, blue.) (C) Membrane components are shown in licorices colored as follows: POPE, blue; POPC, red; SSM, gray; POPS, orange; CHL1, yellow. (D) Periodic boundary conditions implemented in MD simulations do not cause artifactitious IC-EC mixing.

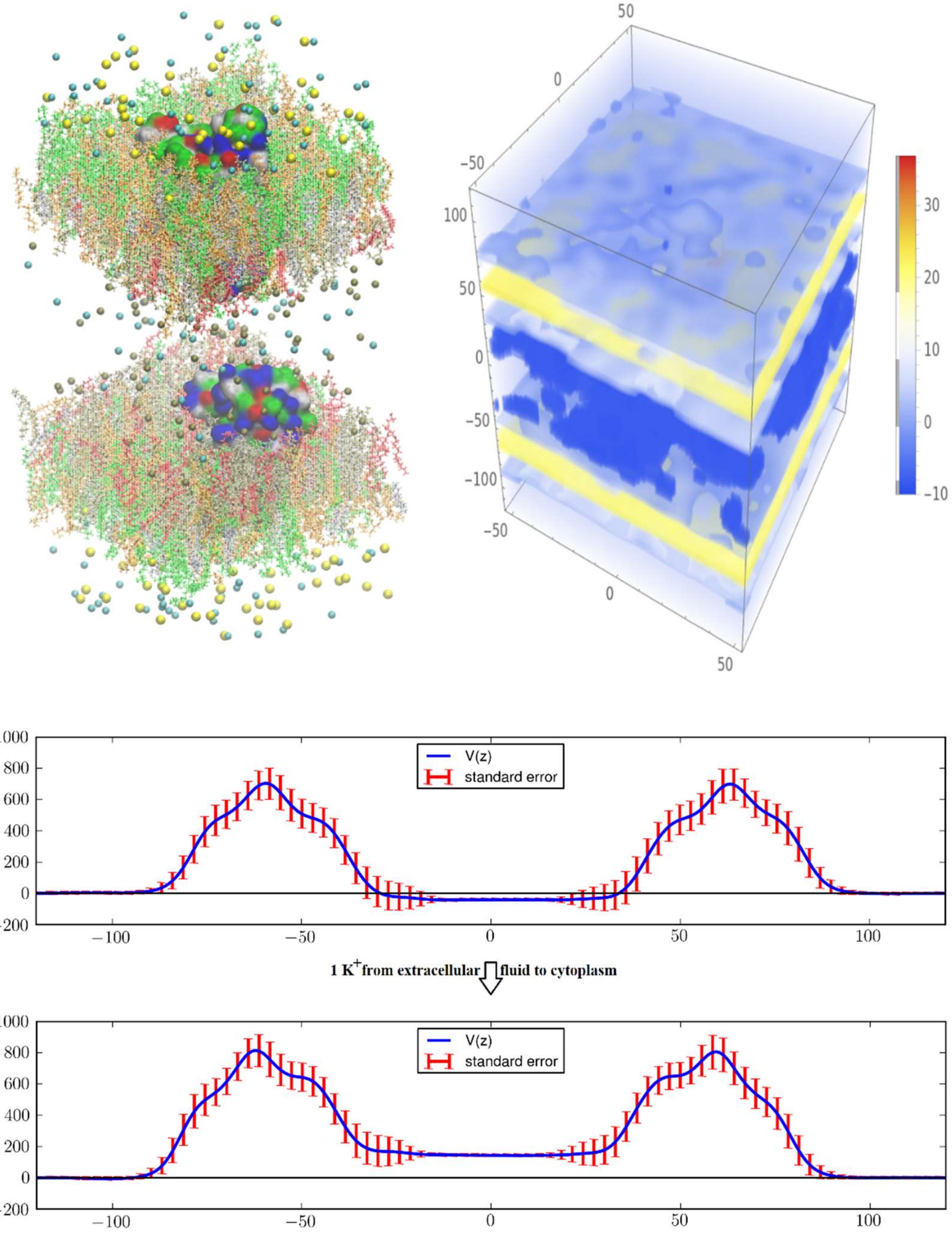


**Fig. S2.** Model system SysI: two GLUT1 proteins in asymmetric environments mimicking the membrane of a human erythrocyte. The intracellular space is located at $-40\text{Å} < z < 40\text{Å}$ and the extracellular space at $z < -95\text{Å}$ and $z > 95\text{Å}$. The water molecules are not shown for clearer views of all the other constituents of the system. The intracellular salt (KCl) and the extracellular salt (NaCl) are shown as spheres colored by atom names ($Cl^-$, cyan; $K^+$, metallic; $Na^+$, yellow). The lipids are represented as licorices colored by lipid names (POPC, orange; POPE, tan; POPS, red; SSM, green; CHL, silver). The proteins are presented as surfaces colored by residue types (hydrophilic, green; hydrophobic, white; positively charged, blue; negatively charged, red). The particle mesh Ewald-based electrostatic potential[4, 5] is shown in the right panel. The membrane potential of this model system is $-30$ mV, which will increase to +168 mV when a single cation is moved across the membrane along the EC-to-IC direction.

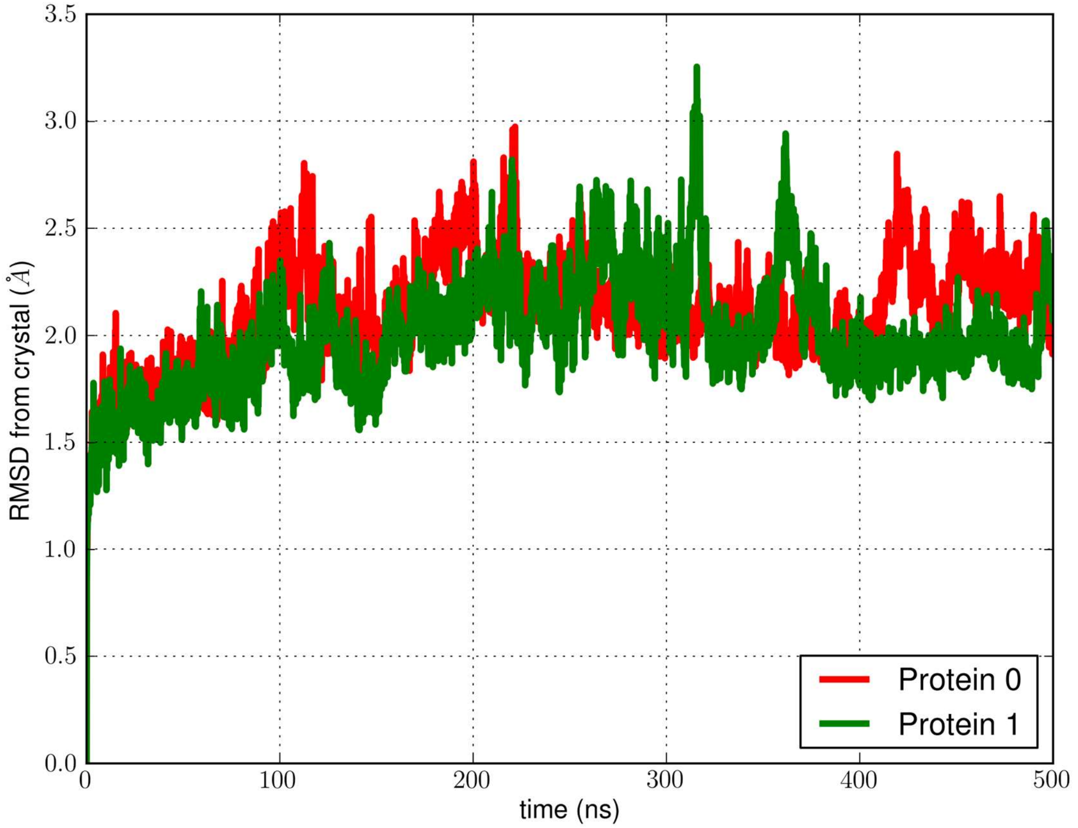


**Fig. S3.** RMSD of GLUT1's in the erythrocyte membrane from the crystal structure during the 500 ns production run of SysI.

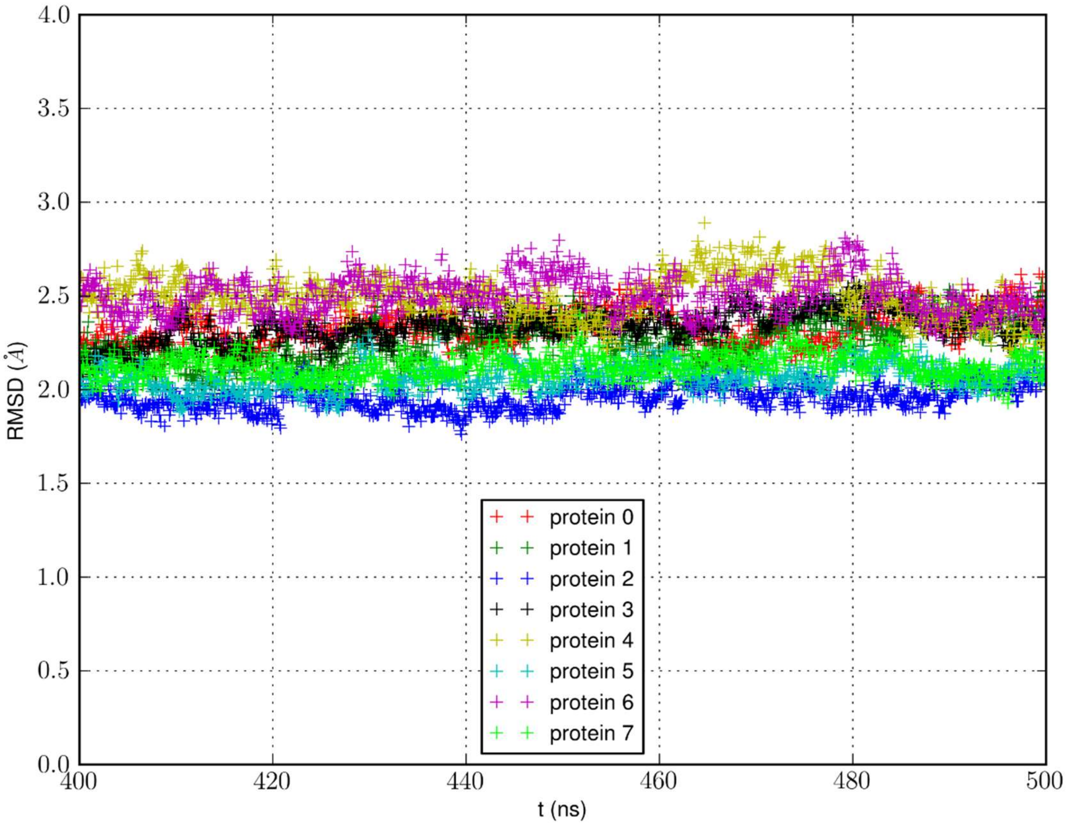


**Fig. S4.** RMSD of GLUT1's in the erythrocyte membrane from the crystal structure during the last 100 ns of the 500 ns production run of SysII, independent repeat 1.

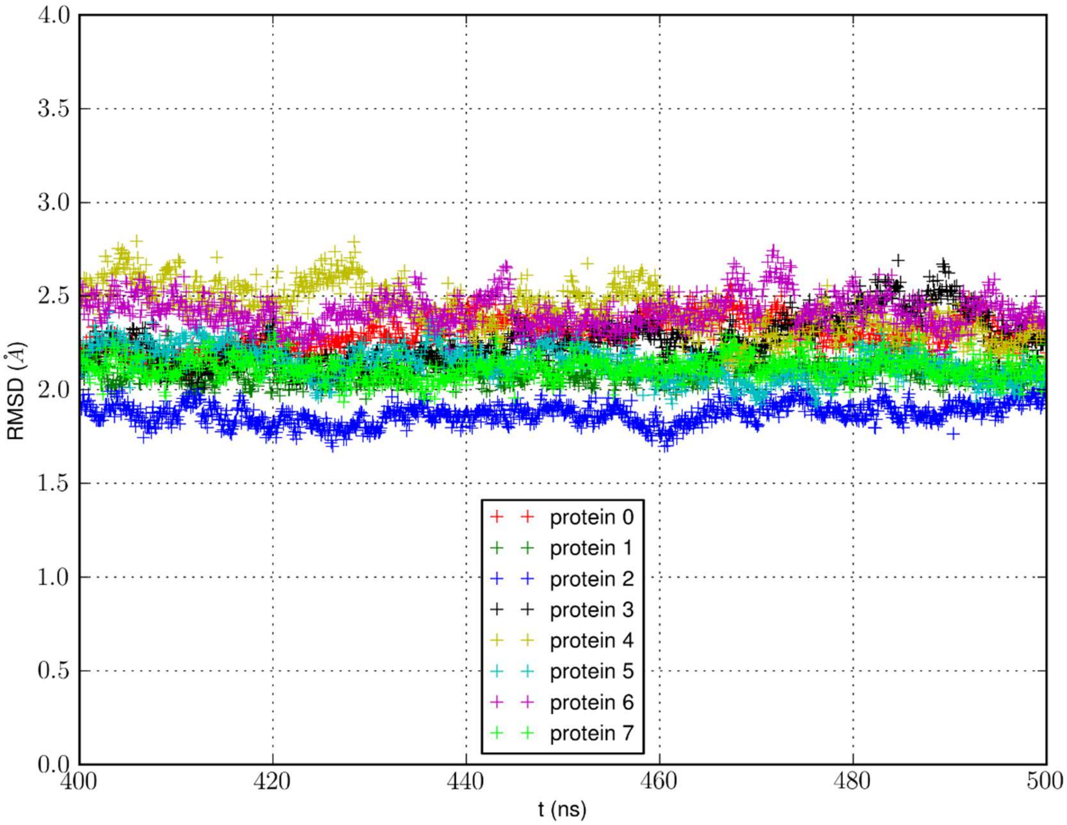


**Fig. S5.** RMSD of GLUT1's in the erythrocyte membrane from the crystal structure during the last 100 ns of the 500 ns production run of SysII, independent repeat 2.

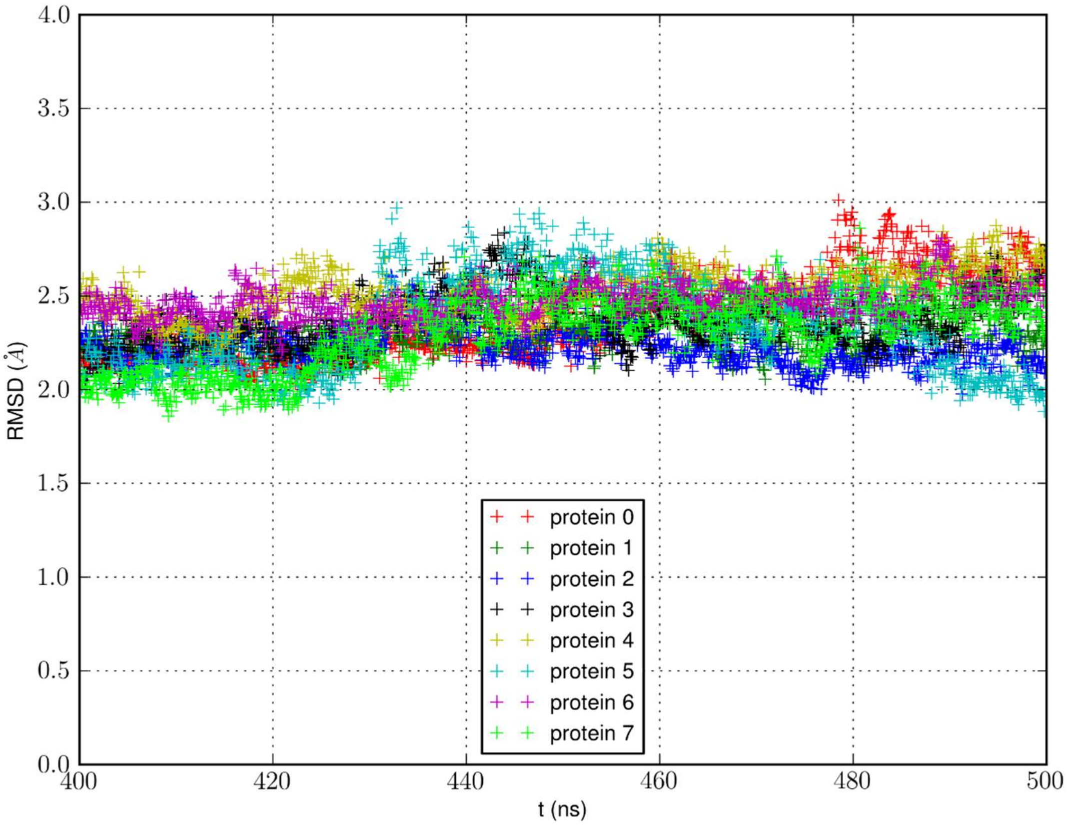


**Fig. S6.** RMSD of GLUT1's in the erythrocyte membrane from the crystal structure during the last 100 ns of the 500 ns production run of SysII, independent repeat 3.
